# Evaluating approaches to find exon chains based on long reads

**DOI:** 10.1101/066241

**Authors:** Anna Kuosmanen, Veli Mäkinen

## Abstract

**Motivation:** Transcript prediction can be modelled as a graph problem where exons are modelled as nodes and reads spanning two or more exons are modelled as exon chains. PacBio third-generation sequencing technology produces significantly longer reads than earlier second-generation sequencing technologies, which gives valuable information about longer exon chains in a graph. However, with the high error rates of third-generation sequencing, aligning long reads correctly around the splice sites is a challenging task. Incorrect alignments lead to spurious nodes and arcs in the graph, which in turn lead to incorrect transcript predictions.

**Results:** We survey several approaches to find the exon chains corresponding to long reads in a splicing graph, and experimentally study the performance of these methods using simulated data to allow for sensitivity / precision analysis. Our experiments show that short reads from second-generation sequencing can be used to significantly improve exon chain correctness either by error-correcting the long reads before splicing graph creation, or by using them to create a splicing graph on which the long read alignments are then projected. We also study the memory and time consumption of various modules, and show that accurate exon chains lead to significantly increased transcript prediction accuracy.

**Availability:** The simulated data and in-house scripts used for this article are available at http://cs.helsinki.fi/u/aekuosma/exon_chain_evaluation_publish.tar.gz.

## 1 Introduction

Third-generation sequencers Pacific Biosciences’ PacBio and Oxford Nanopore increased the sequencing read length tremendously compared to the next-generation platform Illumina. Whereas Illumina read lengths vary from approximately 75 bases to 400 bases, PacBio platform generates reads up to several tens of thousands bases long.

However, due to library preparation limitations, full-length reads for transcripts longer than 2.5 kilobases are less likely to be sequenced (Sharon *et al*., 2013). But even non-full-length long reads can give valuable information about non-adjacent exons in a transcript: Assuming a genome reference is available, one can *align* the long reads allowing introns to be spliced out in the alignment. Then one can read an *exon chain* corresponding to the read alignment, that is, the sequence of exons that the read overlaps.

The prediction of full transcripts can be modelled as a combinatorial problem in a *splicing graph* (Heber *et al*., 2002), where nodes are exons and arcs are exons consecutive in some read alignment. Rizzi *et al*. (2014) proposed modeling long reads as subpath constraints (exon chains) in a splicing graph. Figure 1 illustrates these concepts.

**Figure 1.**
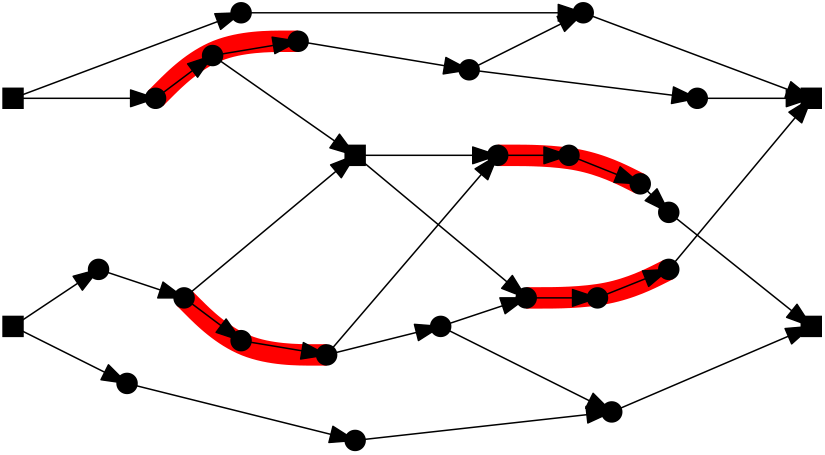
An example of a splicing graph reproduced from Kuosmanen *et al*. (2016), in which three exon chains are drawn in red. The square nodes are the ones where the transcripts can start or end. Exon chains limit the way transcripts could be formed, as each chain should be contained in at least in one transcript.

Kuosmanen *et al*. (2016) implemented the concepts proposed by Rizzi *et al*. (2014) in the transcript prediction tool Traphlor, and using simulated data demonstrated against two state-of-the-art transcript prediction tools StringTie (Pertea *et al*., 2015) and FlipFlop (Bernard *et al*., 2014) that using the information provided by exon chains can significantly improve the transcript prediction accuracy.

However, with the high error rates of the third-generation sequencers (approximately 15% for PacBio and up to 45% for Nanopore, compared to the 1% error rate of second-generation sequencers), aligning the long reads correctly around the splice sites is a difficult task. An additional challenge is posed by the error types: second-generation sequencing errors are generally substitutions, whereas third-generation sequencing errors are mostly insertions and deletions. Incorrect alignments lead to spurious extra nodes and arcs in the splicing graph, which in turn lead to erroneous transcript predictions.

In Figures 2 and 3 we have replicated the experiments of Kuosmanen *et al*. (2016) on a smaller data set simulated from human chromosome 2. As can be seen in the figures, introducing even just mapping errors (as a reminder, in the experimental setup of Kuosmanen *et al*. (2016) no sequencing errors were simulated) causes the accuracy of the prediction to decrease significantly.

**Figure 2.**
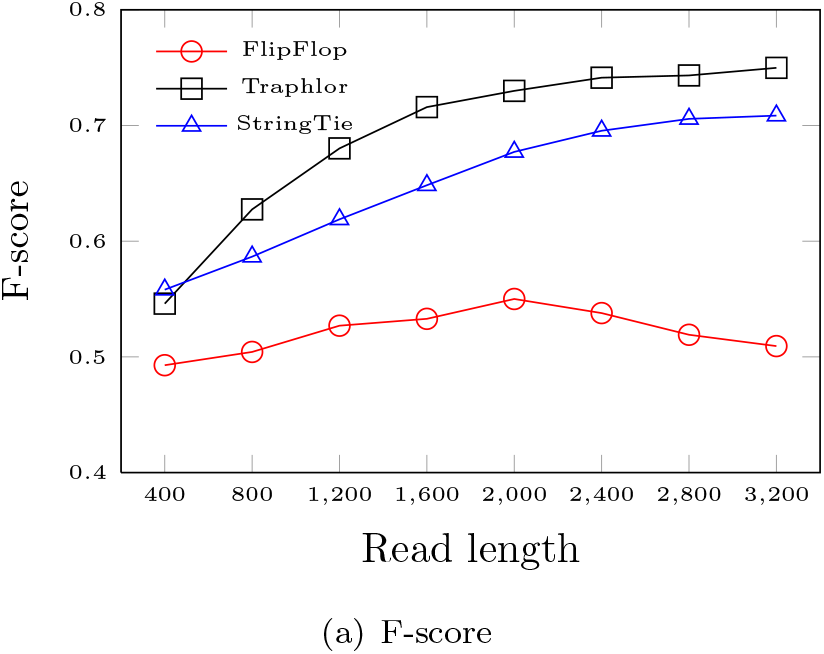
F-score, the harmonic mean of sensitivity and precision, of different transcript prediction tools, StringTie (Pertea *et al*., 2015), FlipFlop (Bernard *et al*., 2014), and Traphlor (Kuosmanen *et al.*, 2016), using *perfect alignments;* alignment information was gathered directly from simulated reads mimicing the use of a perfect aligner.

**Figure 3.**
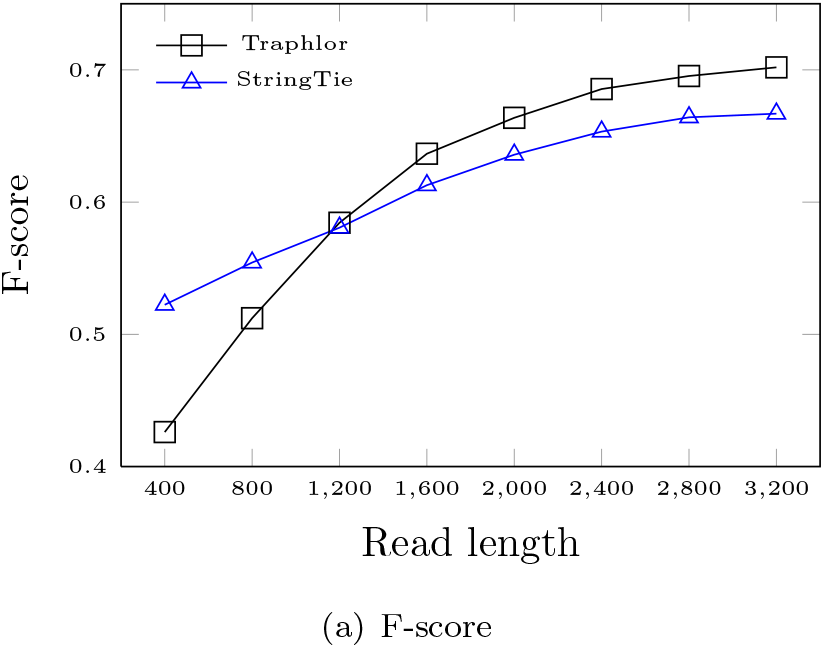
F-score of different transcript prediction tools using GMAP software Wu and Watanabe (2005) for aligning error-free reads.

As suggested in Kuosmanen *et al.* (2016), one approach to tackle the problem of alignment errors near the splice sites would be to optimize the correctness of the chain of exons instead of the alignment of each long read separately.

In this article we survey several approaches to find the exon chains that correspond to the long reads in a splicing graph, as well as provide an experimental study on the performance of these methods. Many of the techniques surveyed are well-known in the literature, but their combination and experimental evaluation towards the identification of exon chains has not been carried out previously. Our study gives several new insights into the feasibility of the problem, and proposes important directions for further studies (see Discussion).

Besides providing new insights, we also obtain results with immediate practical value: We identify practical exon chain identification scenarios to significantly improve the preprocessing steps of transcript prediction tool Traphlor (Kuosmanen *et al*., 2016); it is now feasible to use third-generation long read RNA sequencing to improve transcript prediction accuracy.

## 2 Methods

In this section we introduce three methods that can be used for finding exon chains. As the input we assume there to be both *short reads* and *long reads* from the RNA transcripts. In addition, we assume the reference genome of the species under study to be available, so that we can exploit RNA to DNA read alignments. We consider short reads to be reads of lengths 75–250 with error profile consisting mostly of substitutions, which is typical to the most commonly used second-generation sequencing platform Illumina. Long reads on the other hand are reads with lengths from several hundred to several thousand bases, produced by third-generation sequencing platforms such as PacBio and Oxford Nanopore, and their dominant error type is insertions, followed by deletions. The error rate on the long reads is also a magnitude higher than on the short reads (15-45% vs. 1%).

The first step in finding exon chains is to build *a splicing graph* (Heber *et al*., 2002). In a splicing graph, nodes correspond to exons, and there is an edge between two nodes *v_i_* and *v_j_* if there exists a split read alignment where the exons corresponding to nodes *v_i_* and *v_j_* are consecutive. This is illustrated in Fig. 4.

**Figure 4.**
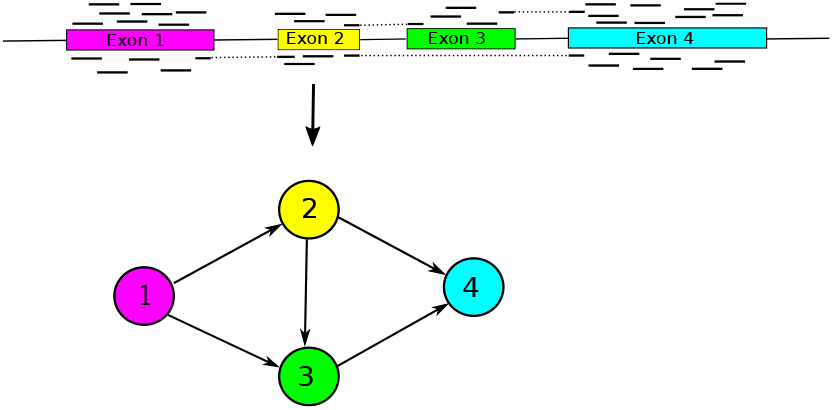
In a splicing graph, exons are represented by nodes, and there is an arc between two nodes if the corresponding exons are consecutive in some read alignment.

Our baseline method is to build the splicing graph using both the short and long reads, and find the exon chains corresponding to the long reads in the resulting splicing graph by matching the genomic coordinates (see Section 2.2 for details). As accurate long read alignment around the splice sites is very difficult (the very fact that prompted this study), using the long reads as-is alone is not feasible.

For comparison, we consider aligning the long reads directly on a splicing graph created from the short reads using dynamic programming (see Section 2.3), as well as a more approximate, but significantly faster, approach of considering the overlaps in genomic coordinates to infer the exon chains (see Section 2.4).

Additionally, we talk about error-correcting the long reads with short reads (see Section 2.1), which can be used both as a pre-processing step as well as a standalone approach. As error-correction significantly increases the mappability of the reads, the alignments around the splice sites are more reliable than when using raw long reads, and the long read alignments alone can be used for inferring the splicing graph.

### 2.1 Error-correcting long reads with short reads

While PacBio reads have very high error rate (> 15%) (Chaisson and Glenn, 2012), and as such pose a harder challenge to error correction than next-generation sequencing reads, the errors seem to be uniformly distributed and independent of the sequence context. Because of this type of error profile, consensus-based methods are suitable for the problem.

There are two main approaches for error-correcting long reads: *self correction* and *hybrid correction*. Self correction uses only long reads, and creates a consensus sequence by computing local alignments between the long reads. Hybrid correction employs more accurate short next-generation sequencing reads to create a consensus by aligning them on the long reads (see Figure 5 for an example).

**Figure 5.**
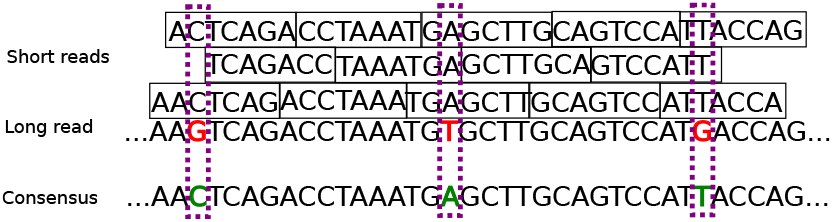
Long reads with high error rate can be corrected by aligning more accurate short reads on them and taking the consensus of the column. In this example the errors in the long read are marked on red, and the corrected bases are marked on green in the consensus sequence.

For this study we use error correction tool LoRDEC (Salmela and Rivals, 2014). LoRDEC is a hybrid correction method that first builds a de Brujin graph (DBG) from short reads, and then uses the DBG to correct erroneous regions within each long read individually. It has been shown to make most of the sequence alignable with percentage of identity >97%, and to do so with significantly lower running time than any previous self-correction tools.

### 2.2 Creating splicing graph from both short and long reads

Our baseline method for this study is to align both short and long reads to the reference genome, and use both of these alignment sets to infer the splicing graph. As mentioned earlier, in a splicing graph exons are represented by nodes, and there is an arc between two nodes if the corresponding exons are consecutive in some read alignment.

The naive approach on locating the exons is to examine the read coverage at every genomic position: positions where the coverage *c* > 0 are designated exonic, and areas where *c* = 0 are designated intronic. Additional information about the exon borders contained within another exon can be found from split read alignments.

Instead of this naive approach, one can use more sophisticated methods for splicing graph creation, using for example software tool SpliceGrapher (Rogers *et al*., 2012).

SpliceGrapher takes as input the read alignment file and gene model annotation that will be used as the base of the graph. From the gene model SpliceGrapher adds all the exon entries as nodes, and sets arcs between them if they are concecutive in some transcript. Then it constructs what the authors call “short-read exons” from clusters of contiguous ungapped alignments. Any short-read exons contained within the exons inferred from the gene model are discarded. Acceptor and donor sites are predicted from spliced alignments, and the short-read exons are extended to the nearest splice site if they don’t already reach it.

It is possible that there exist two exons that overlap in genomic coordinates (but do not have the exactly same start and end coordinates, which would make them the same exon). As a post-processing step for the splicing graph creation, we split any overlapping exons into “pseudoexons” as illustrated in Fig. 6. Unlike exons, pseudoexons cannot have any overlap between them, which simplifies the next steps.

**Figure 6.**
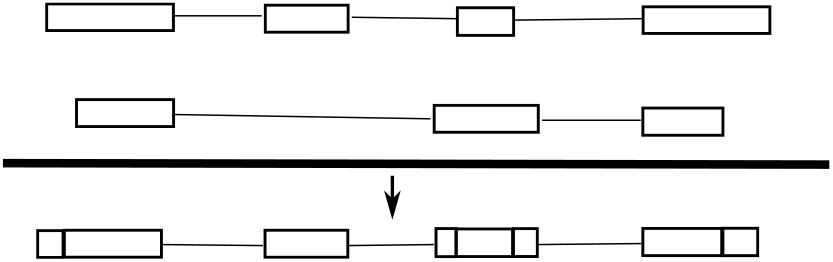
In transcripts two exons can have overlapping genomic coordinates (top). We split the overlapping exons into non-overlapping “pseudoexons” (bottom).

The paths corresponding to the long read alignments can be read from the graph by comparing the start and end coordinates of the blocks (i.e. continuous sequences) in the read alignment to the coordinates of the nodes (pseudoexons). That is, for a node to be considered a candidate in the path, either the start and end coordinates for some block in the alignment have to match the start and end coordinates of the node, or the node has to be completely contained in some block in the alignment. For the first and last block of the read alignment only the end (respectively, start) have to match the coordinates of the node. See Fig. 7 for illustration.

**Figure 7.**
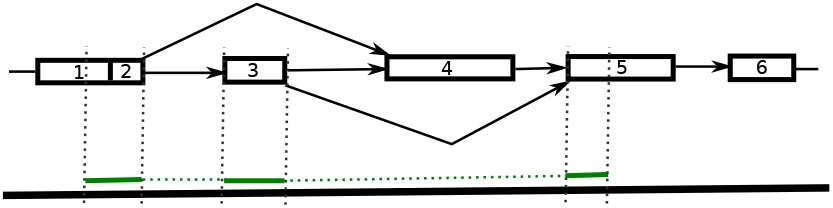
The path corresponding to the long read alignment (green) can be read by comparing the genomic coordinates of the long read alignment and the pseudoexons. In this case the corresponding path is “1, 2, 3, 5”.

If some block of the alignment did not match any node as described above, the alignment is reported to fail to find a path. Also if there is no arc in the splicing graph between two candidate nodes *v_i_* and *v_j_* that are adjacent in the path, the alignment is also reported to fail.

Note that as the long read alignments were also used in the construction of the graph, in theory there should be a path corresponding to every long read alignment. However, this might not be the case in practice if the heuristics of the splicing graph construction algorithm discard some alignments or parts of them.

### 2.3 Aligning long reads to a splicing graph using dynamic programming

We assume a splicing graph has been created using short reads. One can then apply dynamic programming to align a long read into a splicing graph (Mäkinen *et al*., 2015, Section 6.6.5). While this approach is guaranteed to be optimal for the long read in question, other approaches that consider all reads at once may still perform better. (Multiple alignment formulation would give an optimal model, but is infeasible in practice, and will not be discussed here further.)

For self-containedness, we briefly review the approach described in (Mäkinen *et al*., 2015, Section 6.6.5). Denote a directed acyclic graph (DAG) *G* as (*V^G^, E^G^*), where *V^G^* is the set of vertices and *E^G^* is the set of arcs. A *labeled DAG* has a label *ℓ*(*v*) for each vertex *v* ∈ *V^G^*.

*DAG-path alignment problem* is defined as follows. Given two labeled DAGs, *A* = (*V^A^, E^A^*) and *B* = (*V^B^, E^B^*), both with a unique source and sink, find a source-to-sink path *P^A^* in *A*, and a source-to-sink path *P^B^* in *B*, such that the global alignment score *S*(*ℓ*(*P^A^*), *ℓ*(*P^B^*)) is maximum over all such pairs of paths. Here notion *ℓ*() returns the concatenation of vertex labels on the path given as argument.

This problem is easy to solve by extending the global alignment dynamic programming formulation. Consider the recurrence

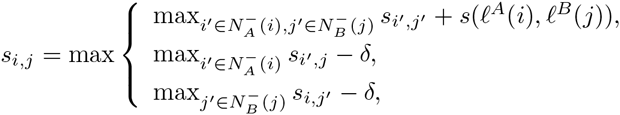

where *i* and *j* denote the vertices of *A* and *B* with their topological ordering number, respectively, 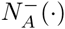 and 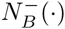 are the functions giving the set of in-neighbors of a given vertex, and *ℓ^A^*(·) and *ℓ_B_*(·) are the functions giving the corresponding vertex labels. Initializing *s*_0,0_ = 0 and evaluating the values in a suitable evaluation order (for example, *i* = 1 to |*V^A^*| and *j* = 1 to |*V^B^*|), one can see that *s*_|*V^A^*|,|*V^B^*|_ evaluates to the solution of the DAG-path alignment: an optimal alignment can be extracted with a traceback. The algorithm takes *O*(|*E^A^*||*E^B^*|) time (Mäkinen *et al*., 2015, Section 6.6.5).

One can now set *A* as a linear labeled DAG denoting the long read and set *B* as the sequence graph corresponding to the splicing graph (see Fig. 8). DAG-path alignment on these inputs gives the best alignment of the long read to the splicing graph. To allow a long read to match a subpath (exon chain) instead of a full transcript, one needs to modify the approach for *semi-local alignment*. This is accomplished by initializing *s*_0,*j*_ = 0 for all *j*. After the dynamic programming, one can look for node *j* with the maximum value *s_m,j_*, where *m* is the length of the long read. Traceback from this node reveals an exon chain containing a best match to the read.

**Figure 8.**
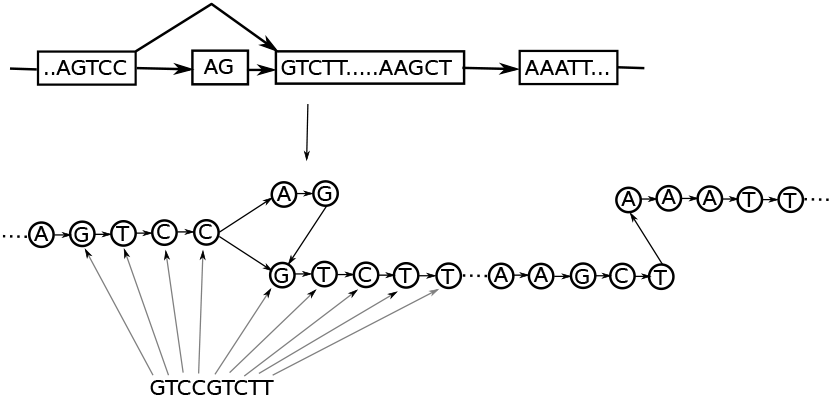
From a splicing graph we create a sequence graph where each base in the sequence is a node. A read sequence can then be aligned to this base graph using dynamic programming.

For the scores for the dynamic programming formula we used the following: match=4, mismatch=-3, insertion=-2 and deletion=-2. This scheme slightly favors insertions and deletions over mismatches, as is appropriate for long read error profile. Choosing lower penalties for insertion and deletion (or higher for substitution) is not possible, as in that case it would be better to insert and delete instead of reporting a mismatch.

### 2.4 Inferring paths in splicing graph from overlaps between exons and aligned long reads

Next we consider a hybrid approach, where we again assume a splicing graph is created using short reads only. But in this case the long reads are aligned to the reference instead of aligning them to the splicing graph. The long read alignments can then be projected to the splicing graphs by examining the coordinate overlaps between the exons in the splicing graphs and the aligned long reads, similarly to the approach described in Section 2.2.

For the purpose of this study, we implemented a naive approach that works as follows. For every long read alignment and for every exon inferred from the short read alignments we calculate the overlap between the coordinates. Every exon *e* that has an overlap with the read *r* is considered a candidate for the path. Then we look for a path *P* = *e*_1_, ⋯ *e_n_*, where the starting coordinates of the candidate exons in the path are in ascending order. If no such path exists in the splicing graph, the path is considered to fail to align.

These concepts are illustrated in Fig. 9.

**Figure 9.**
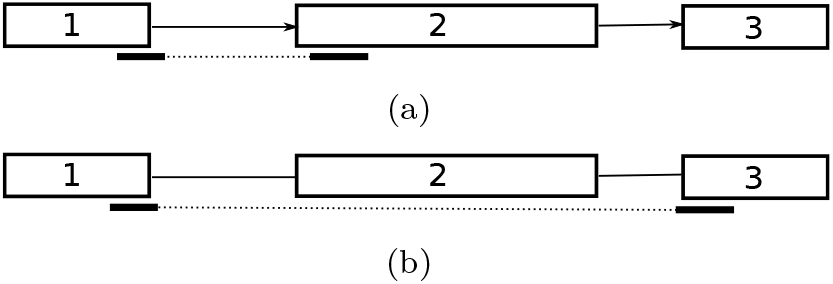
Fig. 9(a): The read overlaps nodes 1 and 2 in genomic coordinates, resulting to potential path of “1 2”. As there is an arc between nodes 1 and 2, the path is accepted. Fig. 9(b): The read overlaps nodes 1 and 3, resulting to potential path of “1 3”. However, there is no arc between the nodes 1 and 3 in the splicing graph, so the path is rejected.

## 3 Results

### 3.1 Data and experimental setup

From human chromosome 2 (version GRCh38/hg38) we sampled all genes that fulfilled the following criteria: 1) shortest transcript was at least 1000 bases long, 2) the longest transcript was at least 3200 bases long, and 3) there were at least two transcripts that did not share all of their inner borders (i.e. used the same exons with different 3’ or 5’ or both). There were a total of 159 genes fulfilling these criteria. We randomly sampled 100 genes for testing.

For every chosen gene, we simulated 10 000 short (75 bp) reads with Illumina-like error profile, consisting of 1% substitution rate, and 1000 long reads of lengths {400,800,1200,1600} with 11% insertion, 4% deletion and 1% substitution rate, which is reported as the error profile of PacBio reads (Chaisson and Glenn, 2012).

As the number of the long reads was low, we considered the transcripts to have uniform expression levels to guarantee that all the transcripts would get sufficiently sampled. The location of the read along the chosen transcript was also sampled from uniform distribution. For read lengths longer than 1000 bases, if the sampled transcript was shorter than the read length, the full-length transcript was added to the data set.

For simulating Illumina-like reads we used RNASeqReadSimulator (Li, 2012). As RNASeqReadSimulator only simulates substitutions, for simulating PacBio-like reads we used an in-house script. The insertions and deletions simulated by the script have length 1.

For comparison we also simulated both short and long reads without any errors.

For creating the splicing graphs we used the software tool SpliceGrapher (Rogers *et al*., 2012) (version 0.2.4). As SpliceGrapher requires being given a gene model in advance, we gave it the annotated transcripts used for the simulation. In addition to the gene model, SpliceGrapher takes as an input a SAM format alignment file. For the alignment of the reads we used GMAP (Wu and Watanabe, 2005) (version 2014-10-22).

For post-processing of the splicing graph we used an in-house script to split any overlapping exons into pseudoexons, as explained in Section 2.2. The script is trivial: all the start and end coordinates of the exons were collected, and based on these coordinates, the pseudoexon list was built.

As described in the previous section, we tested four different approaches: 1) creating a splicing graph from both aligned short and long reads, 2) creating a splicing graph from aligned short reads, and using dynamic programming to align the splicing graph and the long-read-sequences-converted-into-graphs, 3) creating a splicing graph from aligned short reads, and inferring the path by calculating overlaps between the nodes in the splicing graph and the genomic locations of the long read alignments, 4) error correcting the long reads using short reads, aligning only long reads and using these alignments to create a splicing graph.

For cases 1, 2 and 3 we had two subcases, one for reads with simulated sequencing error, and one for reads without simulated error. Additionally, we experimented on using error-correction as preprocessing step and then applying case 2.

To examine how the correctness of the paths affects the downstream analysis, we assembled the transcripts from these data sets using software StringTie (Pertea *et al*., 2015) (version 1.0.1) and Traphlor (Kuosmanen *et al*., 2016). Both StringTie and Traphlor use minimum-cost flows to choose paths in the splicing graph.

StringTie has an integrated mechanism to predict transcripts directly from reads, while Traphlor can make use of the exon chains. The hypothesis is that the better the exon chains are the better Traphlor should perform in comparison to StringTie.

We attempted the downstream analysis also using Cufflinks (Trapnell *et al*., 2010), SLIDE (Li *et al*., 2011a), IsoLasso (Li *et al*., 2011b) and FlipFlop (Bernard *et al*., 2014). But for the bigger read lengths with realistic error profiles we were either unable to run the tools due to memory allocation errors (Cufflinks, FlipFlop) or the tools didn’t produce any output (SLIDE, IsoLasso).

During our experiments we found that SpliceGrapher sometimes creates many erraneous nodes (e.g. a gene model having only 8 nodes resulted in 306 nodes in the predicted graph when using both long and short read alignments). Traphlor’s original problem formulation requires covering all the nodes in the flow network, but due to this problem we relaxed the constraint to only require covering all the nodes corresponding to exon chains.

### 3.2 Validation criteria

For validation we created the “ground truth” paths for all long reads by converting long read BED files created by RNASeqReadSimulator to alignment BAM files using BEDTools (Quinlan and Hall, 2010), and mirroring these alignments on the splicing graph.

For a node to be added to the path, either the start and end coordinates for some block in the alignment had to match the start and end coordinates of the node, or the node had to be completely contained in some block in the alignment. This criteria was relaxed for the first and last block of the read to only require the end (respectively, start) to match the coordinates of the node. For the rest of the article we refer to these paths as “true paths”.

For cases 1 (using both short and long reads) and 4 (error-correcting long reads) we mirrored the aligned long reads on the graph in the same fashion. Cases 2 (calculating overlaps) and 3 (graph alignment) directly produced paths. For the rest of the article we refer to these paths as “predicted paths”.

For all cases, if a sequence of nodes would violate the structure of the graph, in such way that for any two nodes in the path there was no arc between them in the splicing graph, the path was reported to fail.

Two paths were considered to match if they consisted of exactly the same sequence of nodes.

We define *sensitivity* as the number of matched paths divided by the number of true paths, and *precision* as the number of matched paths divided by the number of successfully predicted paths (i.e. excluding the predicted paths that reported to fail). If all the reads successfully produced paths, sensitivity = precision. Otherwise precision can be higher than sensitivity. *F-score*, the standard measure of performance, is the harmonic mean of sensitivity and precision.

For the second part of our experiments, assembling transcripts, we used the same validation criteria as used in (Kuosmanen *et al*., 2016). All predicted transcripts were matched against all the transcripts from which the reads were sampled. Two transcripts consisting of more than one exon were considered a match if all internal boundaries of the transcripts were identical (i.e. the beginning of first exon and the end of last exon did not need to match). Single exon transcripts were considered to match if the overlapping area occupied at least 50% of the length of each transcript. Only one predicted transcript can match a single annotated transcript.

For this experiment, we define *sensitivity* as the number of matched transcripts divided by the number of annotated transcripts and *precision* as the number of matched transcripts divided by the number of predicted transcripts.

### 3.3 Analysis of experiments

As can be seen in the Figure 10, for the data with realistic error profiles both sensitivity and precision were highest using error-correction (case 4) and overlap method (case 3). The best performance was found combining these two methods, which shows that the creation of the splicing graph from the short reads is still slightly more accurate than using error-corrected long reads. As the difference was not significant, we can with high probability assume that the difference was not due to low coverage of the long reads, that is, the coverage was sufficient for the long reads to cover every splicing site.

**Figure 10.**
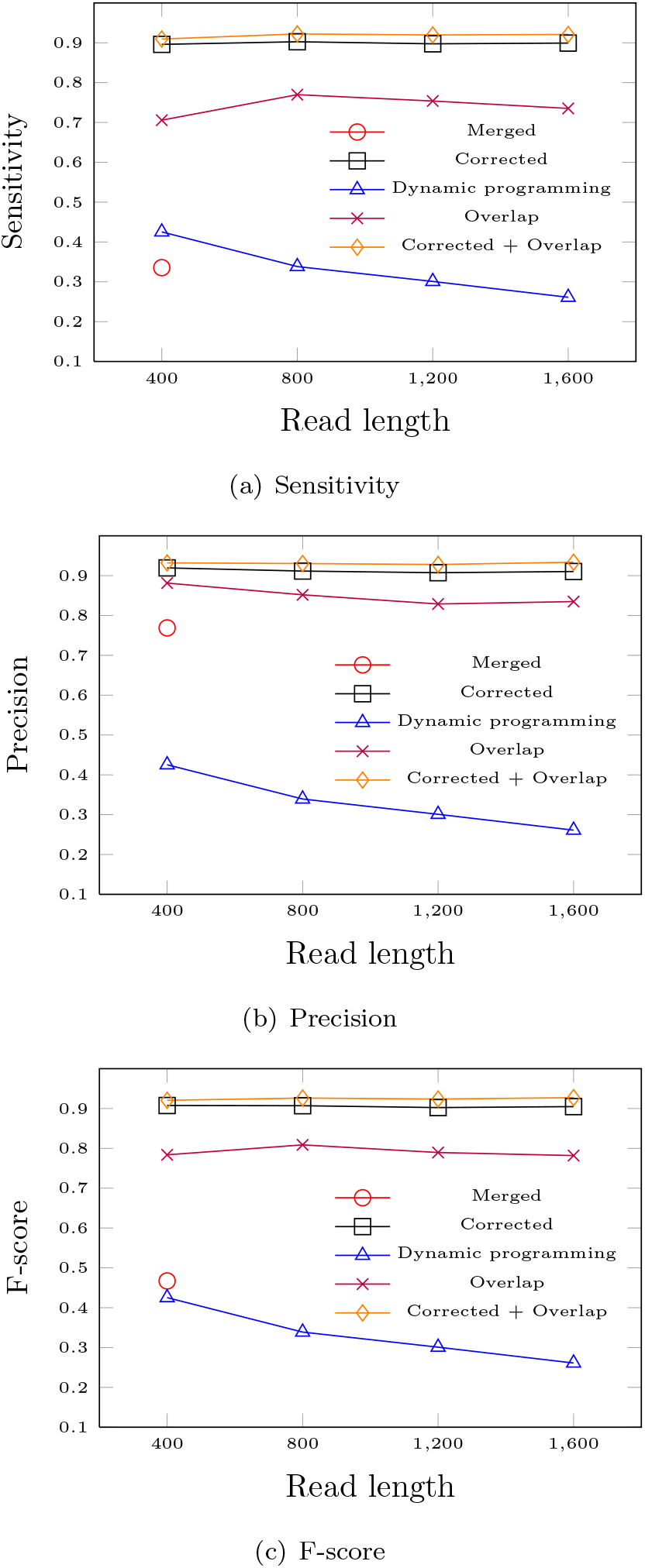
Exon chain prediction accuracy for the cases with 16% sequencing error. In “merged” case long read alignments are mirrored on a graph made from both short and long reads, in “dynamic programming” dynamic programming is used to align long reads on the splicing graph, and in “overlaps” the best overlap in genomic coordinates on the exons predicted from short reads are chosen. In “corrected” the long reads with 16% sequencing error are first error-corrected with short reads, then aligned to the reference and these alignments are mirrored on graph made from short reads. “Corrected + overlap” cases first uses error-correction and then uses the overlaps between short and long read alignments to infer the exon chains.

Surprisingly dynamic programming approach reached only half of the sensitivity and precision of the top performers, and additionally both of the measures decreased as read length increased. While dynamic programming is guaranteed to find some optimal solution for each long read, for read alignment this approach has one severe limitation; if the last base of the exon is the same as the last base of the intron, and both of them are incorporated into the splicing graph, choosing either will give score-wise optimal solution. Whereas for the purpose of validation, only the case that uses the last base of the exon is correct. For the reference, read aligners generally are able to use the information about canonical dinucleotides to break these kind of ties.

Also surprisingly, our baseline method, using both long and short reads in the creation of the splicing graph, had only slightly worse sensitivity and significantly higher precision than dynamic programming approach with read length of 400 bases. As a reminder, precision was defined as the number of matched paths divided by the number of successfully predicted paths. This result is likely due to a large portion of the reads failing to produce paths.

For the cases without errors (see Figure 11), our baseline method achieved sensitivity above 90%, on par with the method using overlaps to infer paths. Dynamic programming achieved approximately 10% higher sensitivity and precision than in the case without errors, but was still significantly worse than the baseline method and the overlap method.

**Figure 11.**
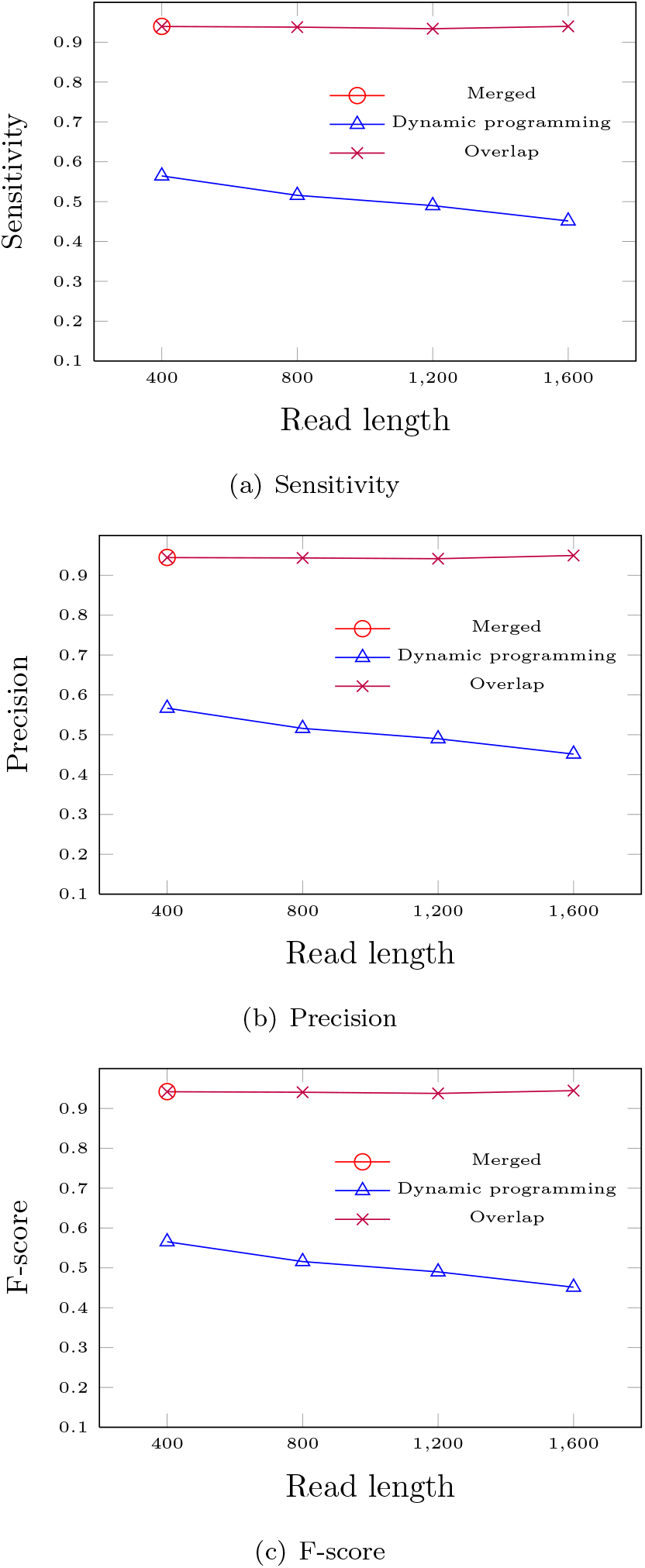
Exon chain prediction accuracy for the cases without sequencing errors. In “merged” case long read alignments are mirrored on a graph made from both short and long reads, in “dynamic programming” dynamic programming is used to align long reads on the splicing graph, and in “overlaps” the best overlap in genomic coordinates on the exons predicted from short reads are chosen.

Due to computational resource constraints, we could only test using both long and short reads in the creation of the splicing graph (the baseline method) for read length of 400 bases. For longer read lengths, the creation of the splicing graph from the highly erroneous long reads took in the excess of 100 hours per gene (i.e. processing the whole data set for one read length would have taken over 13 months).

We also measured the running time (Table 1) and peak memory usage (Table 2) of the different approaches and parts of the pipeline. As expected, the time required for aligning the long reads and error-correcting them increases as read length increases, as does the time required for building the splicing graph from the corrected reads. Excluding the splicing graph creation for both short and long reads, the main bottleneck for the pipeline was the dynamic programming module.

**Table 1.**
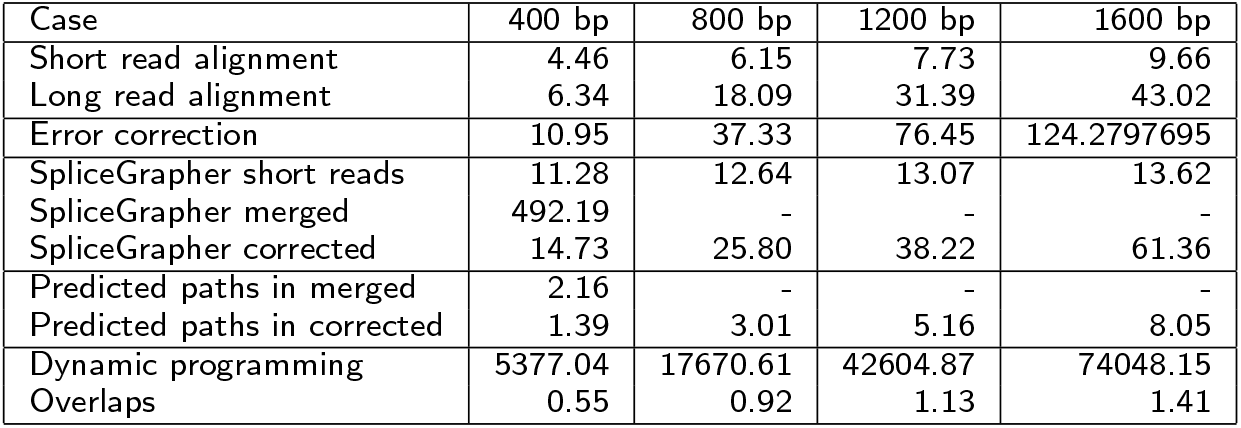
The median running time per gene (in seconds) of different approaches and parts of the pipeline. As the time-measuring module used measured real time instead of CPU time, median time is more suitable than mean time to filter out outliers. For this test we used the data sets with errors only. “Merged” data set includes both short and long reads.

| Case | 400 bp | 800 bp | 1200 bp | 1600 bp |
| --- | --- | --- | --- | --- |
| Short read alignment | 4.46 | 6.15 | 7.73 | 9.66 |
| Long read alignment | 6.34 | 18.09 | 31.39 | 43.02 |
| Error correction | 10.95 | 37.33 | 76.45 | 124.2797695 |
| SpliceGrapher short reads | 11.28 | 12.64 | 13.07 | 13.62 |
| SpliceGrapher merged | 492.19 | - | - | - |
| SpliceGrapher corrected | 14.73 | 25.80 | 38.22 | 61.36 |
| Predicted paths in merged | 2.16 | - | - | - |
| Predicted paths in corrected | 1.39 | 3.01 | 5.16 | 8.05 |
| Dynamic programming | 5377.04 | 17670.61 | 42604.87 | 74048.15 |
| Overlaps | 0.55 | 0.92 | 1.13 | 1.41 |

As memory testing takes several times the normal module execution time, we were unable to test dynamic programming memory requirements for larger read lengths. As the main memory requirement is building the matrix that takes *O*(*nm*) space, where *n* is the length of the reference and *m* is the length of the read, the theoretical peak memory requirement doubles as read length doubles.

While the memory tests were executed on a computing cluster node with 32 GB of RAM, Table 2 shows that all the modules tested fit into 4 GB of RAM, and as such are useable on a regular desktop machine.

**Table 2.** The peak memory usage in megabytes for a randomly picked small subset of the data (n=5), as function of the read length. For this test we used the data sets with errors only. *Dynamic programming values are estimates based on the theoretical bounds. The time for memory testing for them was unrealistic.

| Case | 400 bp | 800 bp | 1200 bp | 1600 bp |
| --- | --- | --- | --- | --- |
| Short read alignment | 653 | 723 | 715 | 664 |
| Long read alignment | 798 | 989 | 1095 | 1145 |
| Error correction | 341 | 341 | 341 | 341 |
| SpliceGrapher (merged alignments) | 3802 | - | - | - |
| SpliceGrapher (short reads) | 3802 | 2538 | 3802 | 2538 |
| SpliceGrapher (corrected reads) | 3802 | 2538 | 3802 | 3802 |
| Find paths (merged alignments) | 10 | - | - | - |
| Find paths (corrected reads) | 10 | 10 | 10 | 10 |
| Dynamic programming | 1184 | 2368* | 3552* | 4736* |
| Overlaps | 10 | 10 | 10 | 10 |

For the downstream analysis we used StringTie and Traphlor (Traphlor base) on the data set containing both short and long read alignments, and additionally gave the splicing graphs and paths created by the various modules to the flow engine of Traphlor. As seen in the Table 3, the running times of both the original Traphlor and Traphlor’s flow engine are linear in read length, but for some reason StringTie’s running time increases by an order of magnitude for every increase in read length. Executing StringTie on 1600 bp was aborted after three CPU days had passed, with the assumption that the process had stalled. However, if the order of magnitude increase keeps up, the expected running time is almost nine CPU days.

**Table 3.** The total CPU time of running the transcript prediction for all 100 genes with long read lengths of 400–1600 bases. When applicable, the time is average between error and no error cases. Note that for StringTie and Traphlor base the time includes creating the graphs. StringTie stalled with 1600 bp, and the merged graph was only created for 400 bp data set.

| Case | 400 bp | 800 bp | 1200 bp | 1600 bp |
| --- | --- | --- | --- | --- |
| StringTie | 16s | 11m 14s | 127m 29s | - |
| Traphlor base | 1m 12s | 3m 8s | 6m 56s | 10m 57s |
| Traphlor merged | 17s | - | - | - |
| Traphlor corrected | 9s | 17s | 21s | 25s |
| Traphlor dp | 11s | 31s | 41s | 1m 14s |
| Traphlor overlap | 4s | 5s | 6s | 6s |

For the transcript predictions based on data with 16% sequencing error, the overall performance of all approaches was low (see Figure 12), with sensitivity in the range of 20-40% for 400 bp reads and 20–60% for the 1600 bp reads, and precision staying below 30%. For the predictions based on the data without sequencing errors (see Figure 13), sensitivity stayed at the same level, but precision for StringTie and for the exon chain approach using overlaps increased significantly, to the range of 40-60%.

**Figure 12.**
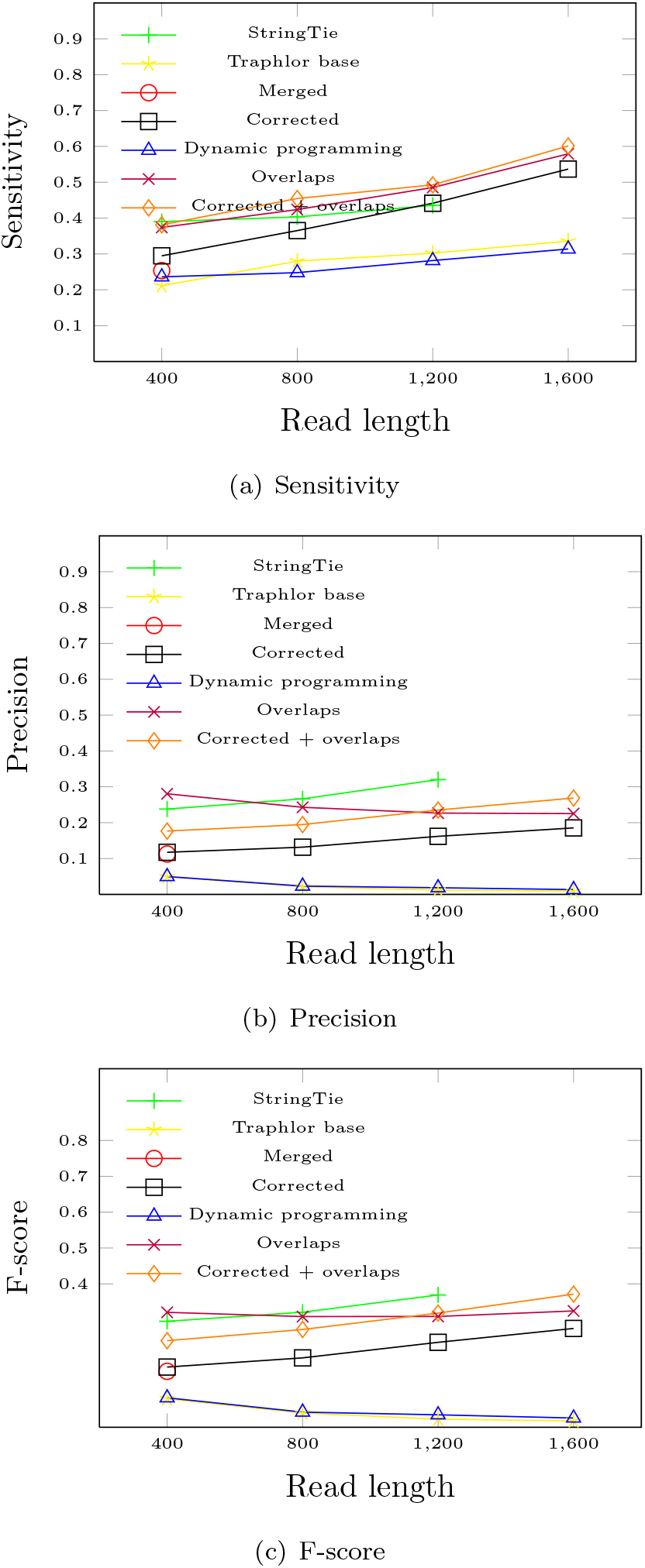
Transcript prediction accuracy using the data sets with 16% sequencing error. Transcripts were predicted from the alignment file (both short and long reads) using software StringTie and Traphlor (Traphlor base), which build their own graphs. In the remaining cases the flow network module of Traphlor was given the predicted graphs and exon chains for each exon chain finding approach.

**Figure 13.**
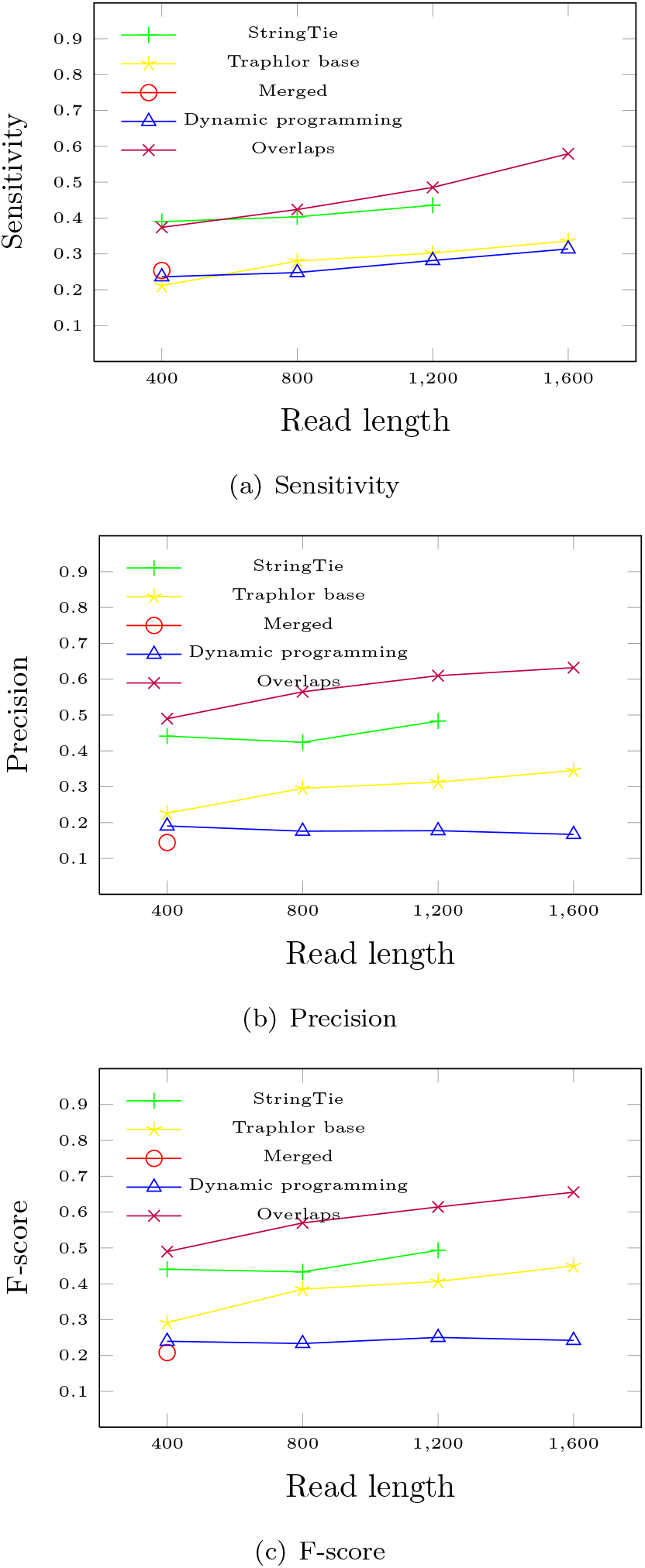
Transcript prediction accuracy using the data sets without sequencing errors. Transcripts were predicted from the alignment file (both short and long reads) using software StringTie and Traphlor (Traphlor base), which build their own graphs. In the remaining cases the flow network module of Traphlor was given the predicted graphs and exon chains for each exon chain finding approach.

Our hypothesis was that the correctness of the exon chains correlates with the accuracy of the transcript prediction, and based on Figures 12 and 13 the hypothesis seems to hold; dynamic programming approach had the worst correctness of the exon chains, and also has the worst transcript prediction accuracy on all three measures. Error-corrected reads alone, overlaps, and error-correction + overlaps approaches have similar performances to each other, with varying rankings between the different measures.

Our baseline methods, Traphlor base and StringTie, ranked as expected. Traphlor base, which contains no heuristics for dealing with errors near splice sites, was systematically either the worst competitor or tied with the dynamic programming approach. StringTie, on the other hand, was among the best. StringTie was however beaten by the overlap approach in sensitivity for both data with and without sequencing errors, and in precision for the data without sequencing errors.

### 3.4 Discussion

In this article we surveyed multiple approaches on finding exon chains, and experimented on the effectiveness of the approaches in finding the exon chains with different read lengths. We also experimented on the effect of the correctness of the exon chains on the downstream analysis (transcript prediction). Additionally, we examined the time and memory requirements of the various parts of the pipeline to identify possible bottlenecks.

For exon chain prediction accuracy, using short reads to correct the long reads was clearly superior both in sensitivity and precision. It is noteworthy that the error correction software that was used is not even tailored for RNA-seq reads, and so the error correction results are likely to improve in the future.

Using short reads to create the splicing graph, and considering the overlaps between the coordinates of the splicing graph and coordinates of the long reads aligned to the reference genome also performed well. As expected, in the tests without simulated error it was on par with the f-score of the error correction method.

Combining error correction and examining the overlaps improved the results slightly over only using error-correction, which points to the graph creation from error-corrected reads not being quite as good as creating it from the short reads.

The hypothesis before conducting this study was that the better the exon chain prediction is the better the transcript prediction will be. Based on our experiments this hypothesis seems to hold; dynamic programming was by far the worst approach for finding the exon chains, and the transcript prediction accuracy mirrors this. However, the overlap method scored slightly worse in the exon chain accuracy than the error-correction method, but produced slightly better transcript prediction.

During the experiments we observed that the splicing graph creation tool we used, SpliceGrapher, did not scale well with increasing read length when given both short and long reads. However, the tool did not have such scaling problems with using the error-corrected long reads. Whether this is a problem with using both short and long reads at the same time, or whether the tool is simply not equipped to deal with the high error rate, is unknown.

Additionally, we found that SpliceGrapher sometimes creates many erraneous nodes. While these low-confidence noded are listed as ‘putative” and could in theory be discarded, discarding all putative nodes could discard real novel exons and splicing events.

Based on our study, we think that for the future work the most important direction is to develop a splicing graph creation tool that is both faster and better able to deal with the error profile of third-generation sequencing data.

While dynamic programming did not show promise in this study, based on the results of using overlaps we also believe an approach combining graph alignment with smart splicing site detection could be feasible, using e.g. co-linear chaining (co-linear chaining on graphs being a topic of another manuscript).

## Acknowledgments

This work has been supported by the Academy of Finland (grant 284598 for Center of Excellence in Cancer Genetics Research).

